# Dramatic changes in mitochondrial substrate use at critically high temperatures: a comparative study using *Drosophila*

**DOI:** 10.1101/2020.11.19.389924

**Authors:** Lisa Bjerregaard Jørgensen, Johannes Overgaard, Florence Hunter-Manseau, Nicolas Pichaud

**Author notes:** Author for correspondence (LBJ). Summary statement: *Drosophila* mitochondrial functions persist at temperatures above organismal heat limits but turn to oxidation of alternative substrates as complex I-supported respiration is impaired.

## Abstract

Ectotherm thermal tolerance is critical to species distribution, but at present the physiological underpinnings of heat tolerance remain poorly understood. Mitochondrial function is perturbed at critically high temperatures in some ectotherms, including insects, suggesting that heat tolerance of these animals is linked to failure of oxidative phosphorylation (OXPHOS) and/or ATP production. To test this hypothesis we measured mitochondrial oxygen consumption rates in six *Drosophila* species with different heat tolerance using high-resolution respirometry. Using a substrate-uncoupler-inhibitor titration protocol we examined specific steps of the electron transport system to study how temperatures below, bracketing and above organismal heat limits affected mitochondrial function and substrate oxidation. At benign temperatures (19 and 30°C), complex I-supported respiration (CI-OXPHOS) was the most significant contributor to maximal OXPHOS. At higher temperatures (34, 38, 42 and 46°C), CI-OXPHOS decreased considerably, ultimately to very low levels at 42 and 46°C. The enzymatic catalytic capacity of complex I was intact across all temperatures and accordingly the decreased CI-OXPHOS is unlikely to be caused directly by hyperthermic denaturation/inactivation of complex I. Despite the reduction in CI-OXPHOS, maximal OXPHOS capacities were maintained in all species, through oxidation of alternative substrates; proline, succinate and, particularly, glycerol-3-phosphate, suggesting important mitochondrial flexibility at temperatures exceeding the organismal heat limit. Interestingly, this compensatory oxidation of alternative substrates occurred at temperatures that tended to correlate with species heat tolerance, such that heat-tolerant species could defend “normal” mitochondrial function at higher temperatures than sensitive species. Future studies should investigate why CI-OXPHOS is perturbed and how this potentially affects ATP production rates.

## Introduction

The body temperature of ectotherms is closely associated with the temperature of their environment. Accordingly, organismal resistance to temperature effects, i.e. thermal tolerance, is an important trait in shaping the biogeographic distribution of ectotherm species, including insects (Addo-Bediako et al., 2000; Kellermann et al., 2012; Sunday et al., 2019). With projections of increasing average temperatures as well as the frequency and intensity of extreme temperature events through climate change (IPCC, 2014), much effort has been put into using characterisation of species heat tolerance to predict global changes in species distribution (Kingsolver et al., 2013; Sunday et al., 2012). Yet, the physiological shortcomings underlying the loss of function and mortality associated with heat stress are still not fully understood for insects (for reviews see Bowler (2018), González-Tokman et al. (2020) and Neven (2000)).

Some physiological and cellular mechanisms often listed as potential contributors to heat mortality in insects and other ectotherms are inactivation and denaturation of proteins, temperature effects on membrane organisation, unaligned temperature sensitivities (Q_10_) of coupled biochemical reactions as well as insufficient oxygen supply in line with mismatched ATP demand and supply (Hochachka and Somero, 2002; Schmidt-Nielsen, 1990). For terrestrial insects there is little evidence to suggest that deficient oxygen supply to the respiring cells is a cause of heat mortality (Klok, 2004; Mölich et al., 2013; Verberk et al., 2015). For example, it is rarely found that moderate hypoxia or hyperoxia alters heat tolerance as would be expected by the OCLTT (oxygen- and capacity-limited thermal tolerance) hypothesis (see Verberk et al. (2015)). However, this general finding does not exclude the possibility that exposure to extreme temperatures challenges mitochondrial function and their ability to produce ATP via the oxidative phosphorylation process (OXPHOS), which is also discussed in a recent review of the literature on mitochondria and ectotherm thermal limits (Chung and Schulte, 2020).

Metabolic demand increases with temperature and to maintain cellular homeostasis the rate of mitochondrial aerobic respiration must keep pace (Blier et al., 2014; Schulte, 2015). Accordingly, thermal sensitivity of mitochondria has been suggested to be important for thermal tolerance and thermal adaptations of mitochondrial functions have been observed in several ectothermic phyla (Chung et al., 2018; Ekström et al., 2017; Fangue et al., 2009; Harada et al., 2019; Havird et al., 2020; Hraoui et al., 2020; Hunter-Manseau et al., 2019; Iftikar et al., 2010; Iftikar et al., 2014; Kake-Guena et al., 2017; Martinez et al., 2016, see also Chung and Schulte, 2020). Most mitochondrial studies addressing the effects of high temperature in ectotherms have focused on aquatic invertebrates or fish, while only a few studies have used insects, even though they comprise > 70% of all animal species (Stork, 2018) and have the most rapidly contracting muscles in nature (Beenakkers et al., 1984; Candy et al., 1997) (but see Chamberlin (2004), Pichaud et al. (2010; 2011; 2012; 2013) and references below for studies on insect mitochondrial function). In insect flight muscle, mitochondrial respiration and ATP turnover may increase hundredfold when transitioning from rest to flight (Davis and Fraenkel, 1940; Krogh and Weis-Fogh, 1951; Weis-Fogh, 1964), and up to 20-fold in *Drosophila* (Chadwick and Gilmour, 1940). Hence, to sustain this intense activity, insect flight muscle metabolism must be extremely flexible. To our knowledge the most comprehensive investigation of the association between insect heat tolerance and mitochondrial function is a series of studies led by Bowler and co-workers on blowflies. Here, the authors described how flight muscle mitochondria isolated from blowflies that had been exposed to sublethal heat stress *in vivo* displayed impaired mitochondrial function (Bowler and Kashmeery, 1981; Davison and Bowler, 1971), and that the organismal recovery from heat exposure (indicated by regained flight ability) was closely associated with the restoration of mitochondrial respiration (Bowler and Kashmeery, 1979; Davison and Bowler, 1971). Similarly, increased organismal heat tolerance induced by a heat shock treatment was found to mitigate damage to mitochondrial function from subsequent sublethal heat stress both *in vivo* and *in vitro* (El-Wadawi and Bowler, 1995). Substantial evidence suggests that mitochondrial oxygen consumption continues beyond the thermal threshold of movement (CT_max_) (Heinrich et al., 2017; Mölich et al., 2013), which is also supported by the blowfly studies, but an important conclusion is that the coupled reactions in mitochondria are challenged around the organismal heat limits (El-Wadawi and Bowler, 1996). Specifically, the latter study on blowfly flight muscle indicated that complex I could be the site of mitochondrial heat damage following a sublethal heat exposure.

In a recent study, we characterised heat tolerance of 11 *Drosophila* species representing a wide array of ecotypes and found pronounced differences in species heat tolerance which was closely related to the temperature of their current distribution (Jørgensen et al., 2019). In the present study we use a subset of this comparative system to ask: 1) whether and how high temperature affects mitochondrial functions in *Drosophila*, and 2) if heat-induced changes in mitochondrial functions are correlated to organismal heat tolerance. Previous studies on mitochondrial function and temperature relations in *Drosophila* have focused on genetic components (mitochondrial haplotypes) and were measured at less stressful high temperatures (up to 28°C) where organismal function is easily maintained (Pichaud et al. (2010; 2011; 2012; 2013)). In contrast, the present study examined effects of high temperature on the electron transport system (ETS) in six species of *Drosophila* representing low, intermediate and high heat tolerance at temperatures approaching and surpassing the lethal limit (19-46°C). This was examined in permeabilized thoraces using high-resolution respirometry to measure multiple steps of the ETS during OXPHOS and non-coupled respiration. To specifically address the role of complex I as the site of heat damage (El-Wadawi and Bowler, 1996), we investigated if complex I-supported OXPHOS diminished at high temperatures, and examined if other components of the ETS compensated under these circumstances, attesting to mitochondrial flexibility during heat stress in *Drosophila*. Finally, we measured *in vitro* activity of mitochondrial enzymes related to complex I substrate oxidation (pyruvate dehydrogenase, citrate synthase and complex I enzymatic activities) to examine if changes in mitochondrial function were directly related to collapse of protein function.

## Materials and methods

### Experimental animals

The present study used six species of *Drosophila* that we previously characterised with respect to heat tolerance (Jørgensen et al., 2019). The species are listed here with increasing level of heat tolerance and the temperature reported to cause knockdown after a 1-hour exposure; *D. immigrans*, Sturtevant 1921 (35.4°C)*; D. subobscura*, Collin 1936 (35.6°C); *D. mercatorum*, Patterson and Wheeler 1942 (37.1°C); *D. melanogaster*, Meigen 1830 (38.3°C); *D. virilis*, Sturtevant 1916 (38.8°C) and *D. mojavensis*, Patterson 1940 (41.2°C). Details on population origin can be found in Table 1 of Jørgensen *et al*. (2019). Flies were kept at Aarhus University (Aarhus, Denmark) for several years before shipping them to the Université de Moncton (Moncton, NB, Canada). Upon reception, flies were acclimated to their previous environmental conditions for about three months prior to the start of experiments (i.e. allowing multiple generations before use). Specifically, flies were maintained at 19°C with a diurnal cycle (12:12 LD) in 35-mL vials with approx. 15 mL oat-based Leeds medium (see Andersen et al. (2015)). Parental flies were moved to a fresh vial every 5-7 days to avoid excessive egg density, and newly eclosed flies were transferred to fresh vials every 2-3 days. Only females 4-8 days post-eclosion were used for experiments.

**Table 1:** OXPHOS coupling efficiency (j≈p) for each species at each temperature reported as mean ± s.e.m. Values of j≈p closer to 1 indicate well-coupled mitochondria while values close to 0 indicate poorly coupled mitochondria, at least at complex I level. As it was not possible to measure the parameters required for calculation of j≈p in unstable preparations, the mean and s.e.m. are based solely on stable preparations (the first number in the parentheses, the second refers to the total number of preparations measured for each species × temperature combination). Within a species, dissimilar letters indicate statistically significant differences in the OXPHOS coupling efficiency between temperatures (one-way ANOVA with Tukey’s post hoc test, p < 0.05). Values of s.e.m. < 0.005 are reported as 0.00 to reduce decimal places. Oxygen coupling efficiencies in bold represent the drop below the “healthy” value 0.8 (linearized transformation of respiratory control ratio RCR of 5, Gnaiger (2014)).

| Temperature<br>(°C) | *D. immigrans* | *D. subobscura* | *D. mercatorum* | *D. melanogaster* | *D. virilis* | *D. mojavensis* |
| --- | --- | --- | --- | --- | --- | --- |
| 19 | 0.92 ± 0.01<br>(8/8)<br>A | 0.93 ± 0.01<br>(7/7)<br>A | 0.98 ± 0.00<br>(8/8)<br>A | 0.90 ± 0.02<br>(7/7)<br>A | 0.95 ± 0.01<br>(7/7)<br>A | 0.97 ± 0.01<br>(7/7)<br>A |
| 30 | 0.95 ± 0.01<br>(7/7)<br>A | 0.96 ± 0.001<br>(7/7)<br>A | 0.97 ± 0.00<br>(7/7)<br>A | 0.89 ± 0.03<br>(8/8)<br>A | 0.96 ± 0.01<br>(7/7)<br>A | 0.97 ± 0.01<br>(7/7)<br>A |
| 34 | 0.94 ± 0.01<br>(7/10)<br>A | 0.95 ± 0.01<br>(2/8)<br>A | 0.92 ± 0.01<br>(5/8)<br>A | 0.92 ± 0.01<br>(7/7)<br>A | 0.97 ± 0.01<br>(7/7)<br>A | 0.96 ± 0.01<br>(7/7)<br>A |
| 38 | **0.78 ± 0.04**<br>(5/9)<br>A | **0.38 ± 0.10**<br>(3/8)<br>B | 0.98 ± NA<br>(1/7)<br>A | 0.87 ± 0.03<br>(7/7)<br>A | 0.94 ± 0.01<br>(3/7)<br>A | 0.90 ± 0.00<br>(3/8)<br>AB |
| 42 | **0.20 ± 0.07**<br>(7/7)<br>B | **0.27 ± 0.07**<br>(8/8)<br>BC | **0.38 ± 0.07**<br>(7/7)<br>B | **0.25 ± 0.12**<br>(8/8)<br>B | **0.43 ± 0.14**<br>(6/7)<br>B | **0.38 ± 0.20**<br>(2/7)<br>BC |
| 46 | **0.01 ± 0.18**<br>(7/7)<br>B | **0.04 ± 0.07**<br>(6/6)<br>C | **0.23 ± 0.07**<br>(7/7)<br>B | **0.06 ± 0.20**<br>(6/6)<br>B | **0.27 ± 0.10**<br>(7/7)<br>B | **0.32 ± 0.15**<br>(6/6)<br>C |

*Note for Table 1: For *D. mercatorum* at 38 °C s.e.m. could not be calculated (NA), as it was only* *possible to calculate $j_{\approx p}$ for a single preparation.*

### Mitochondrial oxygen consumption in permeabilized thoraces

High-resolution respirometry was performed on permeabilized thoraces in the Oxygraph-O2K system (Oroboros Instruments, Innsbruck, Austria) using a protocol for *Drosophila* based on Simard *et al*. (2018). The steps for this protocol are outlined below.

### Preparation of permeabilized thoraces

All steps of the permeabilization protocol were performed on ice. Initially, flies were incapacitated on ice and females were then transferred to a petri dish where the thorax was separated from the head and abdomen. Wings and legs were then removed using a razor blade and a pair of fine-tipped forceps. The number of thoraces required to achieve the target mass of 0.4-1 mg for each Oxygraph chamber was species specific, as size differs between species. For the larger species (*D. immigrans*, *D. subobscura*, *D. mercatorum* and *D. virilis*) two thoraces were used for each chamber, while three thoraces were prepared for the smaller *D. melanogaster* and *D. mojavensis*. Isolated thoraces were immediately transferred to a small Petri dish containing an ice-cold biological preservation solution (BIOPS; 2.77 mM CaK_2_EGTA, 7.23 mM K_2_EGTA, 5.77 mM Na2ATP, 6.56 mM MgCl2, 20 mM taurine, 15 mM Na2phosphocreatine, 20 mM imidazole, 0.5 mM dithiothreitol, 50 mM K-MES, pH 7.1). Thoraces were mechanically permeabilized by delicately poking the tissue with fine-tipped forceps, and the thoraces were then incubated in BIOPS supplemented with 62.5 μg mL^−1^ saponin (prepared daily) for 15 minutes on an orbital shaker (220 rpm) for chemical permeabilization. After 15 minutes, the thoraces were transferred to ice-cold respiration medium (RESPI; 120 mM KCl, 5 mM KH2PO4, 3 mM HEPES, 1 mM MgCl_2_, 1 mM EGTA, adjusted to pH 7.2 then added 0.2 % BSA (w/v)), and incubated for 5 minutes on the orbital shaker (220 rpm) to wash out saponin. Prepared thoraces were gently dry-blotted on a Kimwipe to remove excess RESPI solution and weighed (Secura 225D-1s semi-micro balance (0.01 mg) or Cubis MSE6.6S-000-DM micro balance (0.001 mg), Sartorius, Göttingen, Germany) before they were returned to a droplet of RESPI medium placed on parafilm over ice, such that each RESPI droplet contained the permeabilized thoraces for a single chamber.

### Oxygen consumption rates

Mitochondrial oxygen consumption was measured at six different temperatures: 19°C (acclimation temperature), 30, 34, 38, 42 and 46°C to cover both benign and extreme temperatures for all species. The Oxygraph chambers were set to the assay temperature prior to air calibration, then filled with 2.3 mL RESPI medium and the stoppers were fully inserted to avoid air bubbles. Excess RESPI was aspirated, the stoppers were lifted using the spacer, and the system was allowed at least 45 min with stirring (750 rpm) to equilibrate with the gas phase (air) and stabilise the oxygen concentration dissolved in the medium (solubility decreasing with increasing temperature). When the oxygen signal was stable (as per the recommended ± 1 pmol O_2_ s^−1^ mL^−1^), the system was calibrated relative to the barometric and water vapour pressure (DatLab, Version 6.1.0.7, Oroboros Instruments, Innsbruck, Austria).

Once the Oxygraph had been calibrated, a general substrate-uncoupler-inhibitor titration (SUIT) protocol was employed to measure mitochondrial oxygen consumption at specific steps of the ETS. These steps are described below and are also outlined in Fig. 1, which will be referred to in parentheses throughout the protocol. Concentrations reported here are calculated final concentrations in the 2-mL Oxygraph chamber. Measurements started with removing the chamber stopper and adding 10 mM pyruvate (prepared daily) and 2 mM malate (step 1) followed by the pre-weighed permeabilized thoraces. The oxygen concentration in the chamber was raised to ~150-175 % air-saturation to avoid any oxygen diffusion limitation in the tissue, and the chambers were closed. When oxygen consumption rate was stable, this was taken as the LEAK respiration at the level of complex I (CI-LEAK), which is a non-phosphorylating respiration rate. Injection of 5 mM ADP (step 2) coupled the proton gradient created by electron transfer in complex I to phosphorylation of ADP to ATP (CI-OXPHOS). The integrity of the mitochondrial outer membrane was then examined by injecting 10 μM cytochrome *c* (step 3). A disrupted mitochondrial outer membrane would allow the native cytochrome *c*, which is loosely associated with the exterior of the inner mitochondrial membrane, to escape the intermembrane space and subsequently limit the electron transfer between complex III and complex IV (i.e. limiting oxygen consumption). Accordingly, an injection of cytochrome *c* that results in increased oxygen consumption indicates a compromised outer mitochondrial membrane (likely due to the permeabilization), and preparations where oxygen consumption rate increased more than 15 % were discarded from the analysis (Kuznetsov et al., 2008).

**Fig. 1.**
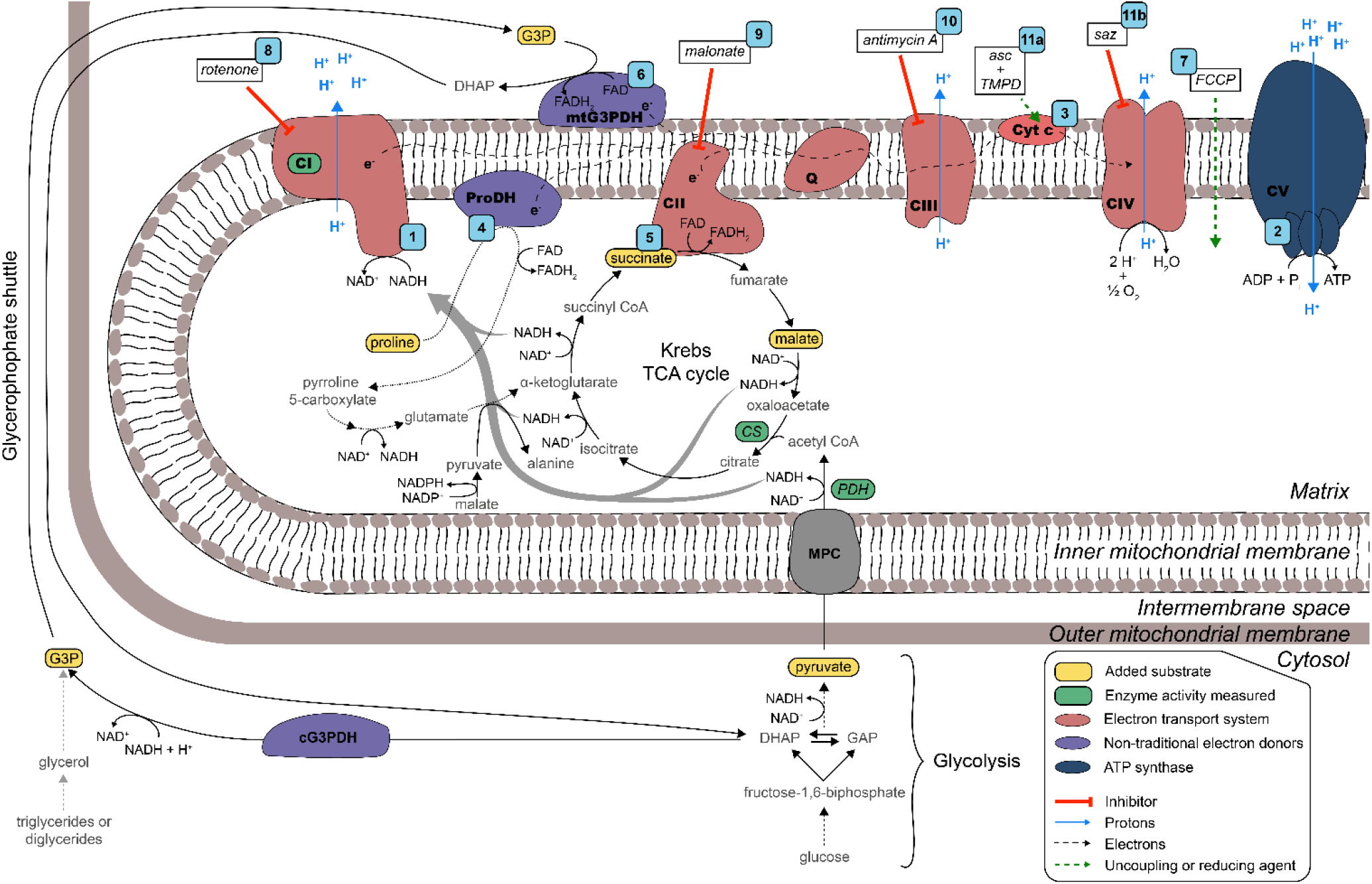
Overview of mitochondrial metabolism and the substrate-uncoupler-inhibitor titration (SUIT) protocol used in the present study. Numbered blue squares refer to steps in the SUIT protocol (also referenced in Materials and methods). In the cytosol, glycolysis transforms glucose into pyruvate which is then transported to the mitochondrial matrix by the mitochondrial pyruvate carrier (MPC). In the matrix, pyruvate dehydrogenase (PDH) transforms pyruvate into acetyl CoA and reduces NAD+ to NADH. Acetyl CoA, which can also be produced by fatty acid oxidation, enters the tricarboxylic acid (TCA) cycle where it participates in condensation with oxaloacetate to form citrate by citrate synthase (CS). The NADH reducing equivalents formed through the TCA cycle and PDH activity are used by complex I (CI) to transfer protons from the matrix to the intermembrane space. Succinate, a TCA cycle intermediate, is oxidized by complex II (CII) which transfers electrons to the ubiquinone (Q) pool via FADH_2_. Proline, a non-traditional electron donor used by some insects, is fueling proline dehydrogenase (ProDH) which transfers electrons directly to the Q pool but can also act as a carbon source and thus replenish the TCA cycle (anaplerotic function, dotted arrows). Glycerol-3-phosphate (G3P), derived from lipid catabolism or transformation of dihydroxyacetone phosphate (DHAP) from glycolysis, is shuttled into the intermembrane space where the mitochondrial glycerol-3-phosphate dehydrogenase (mtG3PDH) reduces the coenzyme FAD and donates electrons to the Q pool. Electrons from upstream complexes in the electron transport system (ETS) converge to the Q pool and subsequently go through complex III (CIII), where protons are pumped to the intermembrane space, and then via cytochrome c (Cyt c) to complex IV (CIV) where molecular oxygen is used as the final electron acceptor and protons are pumped to the intermembrane space. The proton gradient that is formed by the ETS is used by the ATP synthase (complex V, CV) to form ATP through phosphorylation of ADP.

Three additional substrates were added to sequentially stimulate different parts of the ETS. First, 5 mM proline was added as a substrate for proline dehydrogenase (ProDH, step 4) which transfers electrons to the Q-junction in the ETS (CI+ProDH-OXPHOS), followed by succinate (20 mM, step 5), the substrate for complex II (succinate dehydrogenase, CI+ProDH+CII-OXPHOS). Finally, G3P (15 mM, *sn*-glycerol-3-phosphate) was injected (step 6), which is directly oxidized by the mitochondrial glycerol-3-phosphate dehydrogenase (mtG3PDH, CI+ProDH+CII+mtG3PDH-OXPHOS) that similarly feeds electrons to the Q-junction (Fig. 1).

Non-coupled respiration in which the proton gradient produced by the ETS is not coupled to oxidative phosphorylation, and thus indicates the maximal capacity of the ETS, was achieved by titrating the uncoupler FCCP (carbonyl cyanide 4-(trifluoromethoxy)phenylhydrazone, FCCP-ETS) in steps of 0.5-1 μM (step 7). Next the complexes of the ETS were inhibited by injecting 0.5 μM rotenone (complex I inhibitor, step 8), 5 mM malonate (complex II inhibitor, prepared daily, step 9) and 2.5 μM antimycin A (complex III, i.e. blocking the convergent electron transfer from the Q-junction, step 10) to measure the residual oxygen consumption (ROX). ROX was subtracted from all of the substrate-specific oxygen consumption rates to correct for oxygen used by non-mitochondrial oxidative side reactions (see Fig. S1).

The maximal capacity of complex IV (cytochrome *c* oxidase, CIV) for reducing oxygen to water was measured by adding ascorbate (2 mM) and the artificial substrate reducing cytochrome *c*, TMPD (N,N,N’,N,-Tetramethyl-p-phenylenediamine, 0.5 mM) (step 11a). Briefly after the oxygen consumption rate had peaked, it started to decrease due to auto-oxidation of TMPD, and complex IV was immediately inhibited by injection of sodium azide (20 mM, step 11b). The maximal oxygen consumption rate of complex IV was corrected for the underlying auto-oxidation of TMPD by adjusting the slope used to calculate oxygen consumption in DatLab (Version 6.1.0.7, Oroboros Instruments, Innsbruck, Austria).

This SUIT protocol typically took 50-55 minutes, from the time that the permeabilized thoraces were placed in the Oxygraph chambers to the signal stabilization after injection of the last inhibitor (sodium azide). All experiments described above were performed at Université de Moncton.

To examine the temperature sensitivity of electron transport through compartments of the ETS other than complex I, a modified SUIT protocol was applied for additional measurements of oxygen consumption rates at 34 **and 42°C**. Here complex I was blocked with rotenone prior to injection of other substrates, and the CI substrates pyruvate and malate were omitted to minimise reverse electron transport through complex I (Murphy, 2009). Accordingly, the following injections were made (concentrations identical to the general SUIT protocol); rotenone and succinate (CII-LEAK), ADP (CII-OXPHOS), cytochrome *c* (CIIc-OXPHOS), proline (CII+ProDH-OXPHOS), glycerol-3-phosphate (CII+ProDH+mtG3PDH-OXPHOS), FCCP (FCCP-ETS), malonate and antimycin A (ROX, subtracted from the other rates). This SUIT protocol took about 45 minutes until the final injection. The measurements described in this section were performed at Aarhus University, along with additional “control” experiments using the full SUIT protocol described above (not shown). Chemicals were purchased from Millipore-Sigma (Oakville, ON, Canada or Søborg, Denmark).

### Analysis of respiration data

All oxygen consumption rates are here reported as means of mass-specific rates using the unit pmol O_2_ s^−1^ mg^−1^ permeabilized thorax ± s.e.m.

In some experiments, oxygen consumption rate did not stabilise following the addition of ADP (CI-LEAK to CI-OXPHOS transition) but stabilised with the addition of subsequent substrates in the protocol. In these cases, estimates of the CI-OXPHOS oxygen consumption rates were made (see Fig. S1). The unstable traces were observed in five species (not in *D. melanogaster*) and found scattered across temperatures 34, 38 and 42°C. Traces that displayed stable rates during CI-OXPHOS at 34, 38 and 42°C in the five species were analysed to find the average time required for the oxygen consumption rate to stabilise, and the overall mean (138 ± 5 s) across temperatures and species were used to estimate the response to ADP in unstable traces (i.e. the oxygen consumption rate measured 133-143 s after the injection of ADP, see Fig. S1).

CI-LEAK and CI-OXPHOS were used to calculate the OXPHOS coupling efficiency (*j≈P*) at the level of complex I:

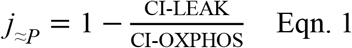

A large increase in oxygen consumption following injection of ADP results in *j≈P* approaching 1, which indicates a highly coupled system as electrons transported by complex I are tightly coupled to oxidative phosphorylation, while an unaffected oxygen consumption rate (*j≈P* = 0) indicates that oxidative phosphorylation does not exert flux control over the electrons transported from complex I (Gnaiger, 2014).

The substrate contribution ratio (SCR), i.e. the relative contribution to increased oxygen consumption rate when adding a new substrate (proline, then succinate followed by G3P) was calculated as

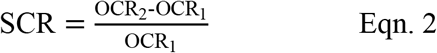

Where OCRM_1_ is the oxygen consumption rate prior to injection of the new substrate (e.g. CI+ProDH-OXPHOS) and OCRM_2_ is the oxygen consumption rate with the new substrate injected (e.g. CI+ProDH+CII-OXPHOS). A value of SCR close to 0 indicates that the added substrate did not increase the oxygen consumption markedly, while SCR = 1 indicates a 100 % increase (doubling), SCR = 2 a 200 % increase (tripling), and so forth.

The maximal ETS capacity where electron transfer is not coupled to phosphorylation (FCCP-ETS) was compared to the maximal oxygen consumption rate coupled to phosphorylation (CI+ProDH+CII+mtG3PDH-OXPHOS) to calculate the non-coupled ratio, ETS_max_/OXPHOS_max_:

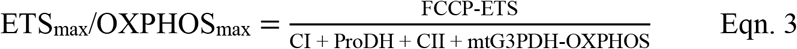

If ETS_max_/OXPHOS_max_ = 1, then the ETS is already fully coupled to phosphorylation, while ETS_max_/OXPHOS_max_ > 1 indicates that the ETS is limited by the downstream process of phosphorylation, and thus have capacity to increase electron transport when the limiting step is alleviated by adding the uncoupler FCCP.

### Enzymatic activities

Measurement of enzymatic activities for all species were performed at similar temperatures as the measurements of mitochondrial oxygen consumption (23.5, 30, 34, 38, 42 and 45°C; note the system could not be cooled to 19°C (substituted by 23.5°C) nor heated to 46°C (substituted by 45°C)). For each species, 6 pools of female flies (4-7 days post-eclosion) were used (N = 6, 10 flies in each pool for all species except the large *D. immigrans* where only 5 flies were pooled), and all measurements were run with 2-3 technical replicates from each pool. Flies were chilled and their thoraces were dissected and stored at −80°C, until they were homogenized in phosphate-buffered saline (137 mM NaCl, 2.7 mM KCl, 10 mM Na_2_HPO_4_, 1.8 mM KH_2_PO_4_, pH 7.4) using a pellet pestle and the resulting homogenates were centrifuged at 750*g* for 5 min at 4°C. The supernatant was then directly used for measurements of NADH:ubiquinone oxidoreductase (complex I, CI) and pyruvate dehydrogenase (PDH). The remaining supernatant was kept at −80°C for later measurement of citrate synthase (CS) and total protein content. All enzymatic activities were measured following protocols already established (Ekström et al., 2017; Cormier et al., 2019) using a BioTek Synergy H1 microplate reader (BioTek®, Montreal, QC, Canada). Enzymatic activities (EA) were calculated using the following equation:

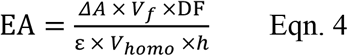

Where ΔA represents the variation in absorbance, V_f_ is the final volume in the well, DF represents the dilution factor, ε is the molar extinction coefficient, V_homo_ represents the volume of homogenate used and h is the height of the volume in the well (including the bottom thickness). The height h is calculated as h = (4V)/(πd^2^) where V is the final volume of the reaction (between 200-225 μL of reaction medium depending on the enzyme measured) and d is the diameter of the well. Concentrations for each compound of stock solutions used are described below:

CI (EC 7.1.1.2) activity was measured by following the reduction of 2,6-dichloroindophenol (DCPIP) at 600 nm (ε=19.1 mL cm^−1^ μmol^−1^). Briefly, CI oxidizes NADH and the electrons produced reduce the ubiquinone 1 (UQ1) which subsequently delivers the electrons to DCPIP. After incubation of homogenates in a 100 mM potassium phosphate buffer containing 0.5 mM EDTA, 3 mg mL^−1^ BSA, 1 mM MgCl_2_, 2 mM KCN, 4.2 μM antimycin A, 75 μM DCPIP and 65 μM UQ1, pH 7.5, for 5 minutes in the plate reader at assay temperature, 0.14 mM NADH was added to start the reaction, which was recorded for 10 min. The same reaction with 1 μM rotenone was followed in parallel and the specific CI activity represented by the rotenone-sensitive activity was calculated.

PDH (EC 1.2.4.1) activity was measured using the reduction of p-iodonitrotetrazolium violet (INT) at 490 nm (ε = 15.9 mL cm^−1^ μmol^−1^) for 10 minutes after homogenates had been incubated in 50 mM tris-HCl, 0.1%(v/v) triton-X100, 1 mM MgCl_2_ and 1 mg mL^−1^ BSA complemented with 2.5 mM NAD, 0.5 mM EDTA, 0.1 mM coenzyme A, 0.1 mM oxalate, 0.6 mM INT, 6 U mL^−1^ lipoamide dehydrogenase, 0.2 mM thiamine pyrophosphate and initiated with 5 mM pyruvate, pH 7.8.

CS (EC 4.1.3.7) activity was determined at 412 nm for 5 minutes by measuring the reduction of 5,5-dithiobis-2-nitrobenzoic acid (DTNB, ε=14.15 mL cm^−1^ μmol^−1^, Riddles et al. (1979)) using a 100 mM imidazole-HCl buffer. containing 0.1 mM DTNB, 0.1 mM acetyl-CoA and 0.15 mM oxaloacetic acid, pH 8.0.

Total protein content was measured using the bicinchoninic acid method (Smith et al., 1985) and subsequently enzymatic activities are reported as U g^−1^ protein, where U represents 1 μmol of substrate transformed to product in one minute.

### Statistics

Statistical data analyses were performed in *R* version 3.6.2 (R Core Team, 2019). For comparison of oxygen consumption rates, calculated ratios and enzyme activities statistical analyses were performed across temperatures within substrate combinations (or ratio type or enzyme) within species using one-way ANOVAs and Tukey’s *post hoc* test using the *emmeans*-function (estimated marginal means) in the emmeans-package in *R* (Lenth, 2019).

Oxygen consumption rates from stable and unstable traces (deemed after ADP injection) were compared using Welch two-sample *t*-tests (see Fig. S1 and accompanying text).

## Results

### High temperature results in loss of mitochondrial complex I oxidative capacity, but maximal respiration is partially rescued by oxidation of alternative substrates

To examine the sensitivity of mitochondrial function to high temperature, mass-specific oxygen consumption rates (OCRs) were measured at six temperatures (19, 30, 34, 38, 42 and 46°C) in permeabilized thoraces from six *Drosophila* species over multiple steps of the electron transport system (ETS). For each step, OCRs were evaluated *between* assay temperatures *within* species with temperature as the fixed factorial variable using one-way ANOVAs and, when applicable, pairwise comparisons with Tukey adjustment of *p*-values (*F*-values: Table S1). To simplify the graphical presentation of the results, three species (*D. immigrans*, *D. mercatorum* and *D. virilis*) are shown in the Supplementary Information (Figs. S2, S3), and thus only measurements from *D. subobscura*, *D. melanogaster* and *D. mojavensis* are presented graphically below. The omitted species have organismal heat tolerances that approximately corresponds to that of the presented species in the order above (e.g. *D. immigrans* and *D. subobscura* have similar (low) heat tolerance).

LEAK-state respiration was assessed at the level of complex I (CI-LEAK) by injecting pyruvate and malate, and this corresponds to the oxygen consumption required to offset the proton leak across the inner mitochondrial membrane from the intermembrane space without phosphorylation. CI-LEAK was generally low in all species across temperatures. However, assay temperature was found to affect the rate in species *D. immigrans* (Fig. S2A), *D. subobscura* (Fig. 2A), *D. mercatorum* (Fig. S2B) and *D. melanogaster* (Fig. 2B), though in the latter the *post hoc* test failed to separate the temperatures. No statistically significant effects of assay temperature were found in *D. virilis* (Fig. S2C) or *D. mojavensis* (Fig. 2C).

**Fig. 2.**
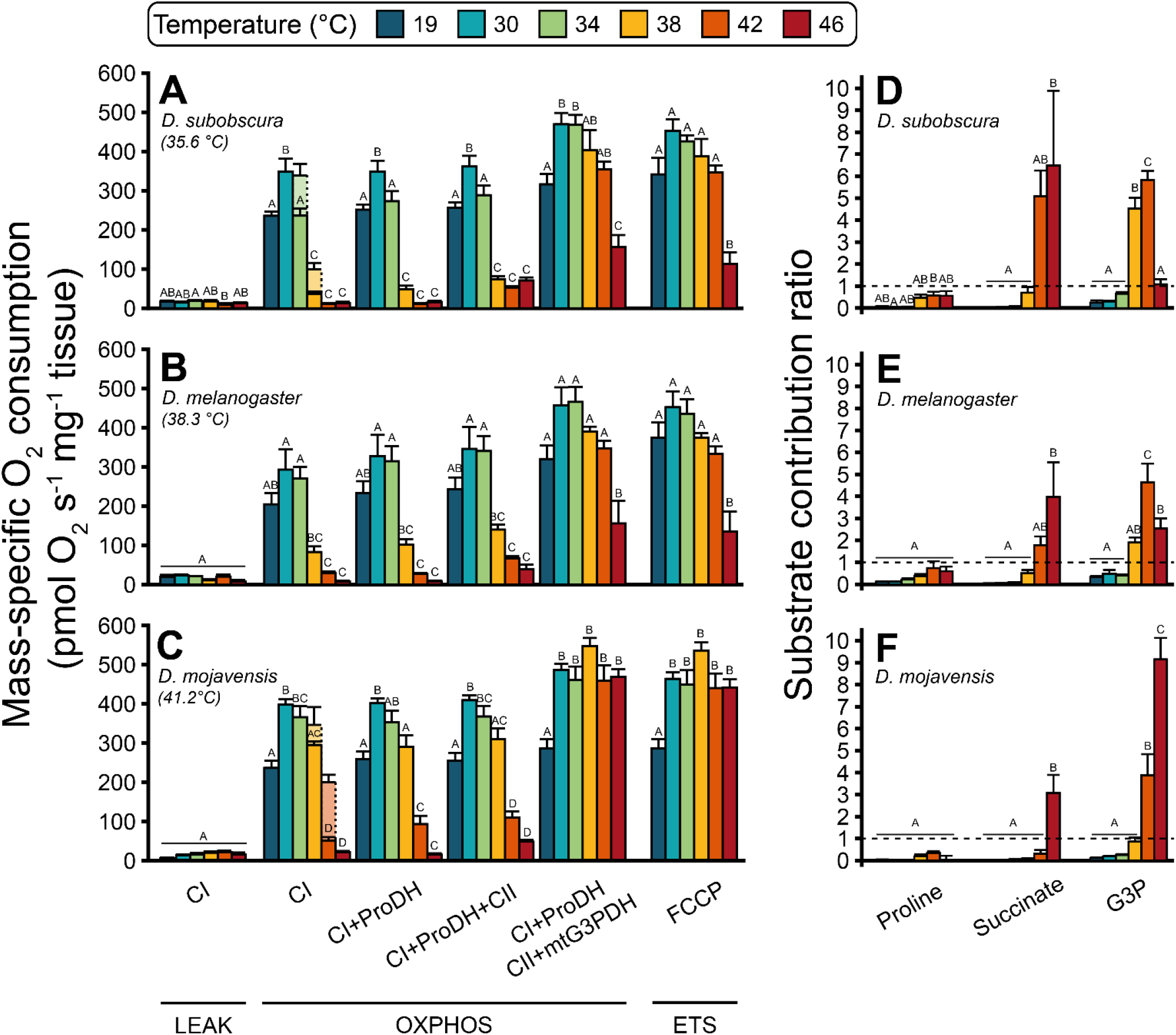
Mass-specific oxygen consumption rates (OCRs) in permeabilized Drosophila thoraces and calculated substrate contribution ratios for three of the tested species. (A-C) Using a substrate-uncoupler-inhibitor titration (SUIT) protocol (Fig. 1), OCRs were measured in the LEAK (without ADP), OXPHOS (oxidative phosphorylation, respiration coupled to phosphorylation with saturating ADP) and ETS (electron transport system, non-coupled from phosphorylation) states. Briefly, complex I substrates pyruvate and malate were provided (CI-LEAK), ADP was added to couple respiration to phosphorylation (CI-OXPHOS), and the OXPHOS state was further measured using successive injections of substrates stimulating different parts of the ETS: proline (CI+ProDH-OXPHOS), succinate (CI+ProDH+CII-OXPHOS) and glycerol-3-phosphate (CI+ProDH+CII+mtG3PDH-OXPHOS). Maximal capacity of the ETS with convergent electron transfer from the pathways above, was measured using the artificial uncoupler FCCP (FCCP-ETS). The SUIT protocol was performed at six assay temperatures, 19°C (maintenance temperature, dark blue) and 30, 34, 38, 42 and 46°C (temperature increasing going towards the rightmost bars in the clusters) covering benign and stressfully high temperatures for the Drosophila species tested (ordered A through C indicating increasing organismal heat tolerance, with temperature estimated to cause knockdown after one hour in parentheses (Jørgensen et al., 2019)). OCRs are reported as mean pmol O_2_ s^−1^ mg^−1^ tissue ± s.e.m., sample size for each species × temperature combination is described in Table 1. In some preparations, oxygen flux did not stabilise following the injection of ADP (CI-OXPHOS) and instead an estimate of the flux was made (lighter shaded, dashed lined bars, see main text and Fig. S1). Within each step of the protocol (cluster of bars), the OCRs were compared between temperatures within species using a one-way ANOVA with a Tukey’s post hoc test, and dissimilar letters within a cluster indicate statistically significant differences (p < 0.05). (D-F) Substrate contribution ratio (SCR) for the three sequentially injected substrates; proline, succinate and glycerol-3-phosphate (G3P). SCR is calculated as the ratio between the increase in OCR following injection of the new substrate compared to the prior OCR, and as such the SCR of proline is based on the change compared to CI-OXPHOS, SCR of succinate on CI+ProDH-OXPHOS and SCR of G3P on CI+ProDH+CII-OXPHOS, respectively. The scale of the SCRs refers to fold changes from the base rate, i.e. 0 means that the added substrate did not change the OCR, 1 refers to a doubling (100% increase from base rate), 2 to three-fold change (200%), etc. SCRs are reported as mean ± s.e.m., sample size for each species × temperature combination is described in Table 1. For preparations that were unstable at CI-OXPHOS it was not possible to calculate the SCR for proline, and accordingly values of SCR for proline are based solely on stable CI-OXPHOS preparations, while SCR for succinate and G3P were calculated for all preparations. Within each species, SCRs were compared between temperatures using a one-way ANOVA with a Tukey’s post hoc test, and dissimilar letters within a cluster indicate significant differences (p < 0.05). In D. melanogaster a significant effect of temperature was found for proline (F5,37 = 2.571, p = 0.043), but the post hoc test failed to reveal significant contrasts between temperatures. The results from the other three species are presented in Fig. S2.

Next, complex I respiration was measured by injecting ADP to couple electron transport (and hence oxygen consumption) to phosphorylation (CI-OXPHOS). Assay temperature was found to affect CI-OXPHOS in all six species (one-way ANOVAs, *p* < 0.001), and there was a general pattern of how this temperature effect was manifested. Increasing temperature from the acclimation temperature (19°C) to 30°C increased CI-OXPHOS for all species (Figs. 2A-C, S2A-C), although for *D. immigrans* and *D. melanogaster* this increase was not statistically significant (Tukey’s *post hoc* adjustment: *p* = 0.177 and *p* = 0.281, respectively). In measurements performed at 34, 38 and 42°C, we observed several unstable preparations with distinctive features; a sharp increase in OCR following ADP injection which was quickly followed by a gradual, consistent decrease that persisted until subsequent substrate injections which led to stabilisation of the OCR (see Fig. S1). The analysis and quantification of these unstable traces is discussed in the paragraph below. At 34°C, stable CI-OXPHOS rates were similar to those measured at 30°C (*D. immigrans*, *D. melanogaster*, *D. virilis* and *D. mojavensis*), while it decreased in *D. subobscura* and *D. mercatorum*, albeit not significantly in the latter species. At 38°C, CI-OXPHOS decreased significantly compared to 34°C in the four least heat tolerant species, i.e. *D. immigrans*, *D. subobscura*, *D. mercatorum* and *D. melanogaster* (Figs. 2A,B, S2A,B). The heat tolerant *D. virilis* and *D. mojavensis* also showed a trend for decreased CI-OXPHOS at 38°C, but the rates were not significantly different from those measured at 34°C (Figs. 2C, S2C). At temperatures above 38°C, all species showed significantly decreased CI-OXPHOS, with non-significant differences between rates measured at 42 and 46°C, although the mean rates were almost always lower at 46°C. Accordingly, all species had CI-OXPHOS rates at the highest temperatures that were significantly lower than at their acclimation temperature (19°C).

As stated above, unstable CI-OXPHOS measurements were found in five of the six species (not *D. melanogaster*), and distributed such that *D. immigrans*, *D. subobscura* and *D. mercatorum* (heat sensitive) displayed instability at 34 and 38°C, while unstable CI-OXPHOS traces were found at 38 and 42°C in *D. virilis* and *D. mojavensis* (heat tolerant) (Table 1). To quantify this observation, we obtained the OCR following ADP injection at the time where “stable” traces would have stabilised CI-OXPHOS (138 ± 5 s) to generate a proxy for CI-OXPHOS in the unstable preparations. The mean of these “unstable” rates was generally higher than the mean of stable CI-OXPHOS at the same temperature within a species (compare shaded and full bars in Figs. 2, S2, see also Fig. S1) which could indicate that the unstable and declining traces were still underway in reaching a lower and potentially stable rate (see discussion).

Following measurements of CI-OXPHOS, cytochrome *c* was injected to test the condition of the outer mitochondrial membrane, and a reduced response (< 15% increase in OCR) was taken as an indication that the outer mitochondrial membrane was intact following permeabilization. Then proline (CI+ProDH-OXPHOS) and succinate (CI+ProDH+CII-OXPHOS) were injected, and the general patterns for the thermal sensitivity of the resulting OCRs were similar to the patterns observed for CI-OXPHOS (similar lettering within species in Figs. 2A-C, S2A-C).

Injection of glycerol-3-phosphate (G3P) gave rise to the maximal OCR during the OXPHOS state, allowing to evaluate the convergent electron flow into the ETS in a coupled state. The OCRs measured as CI+ProDH+CII+mtG3PDH-OXPHOS may be driven by different contributions of mitochondrial complexes and dehydrogenases, and it is therefore of interest to examine the rates both as a measure of maximal coupled respiratory capacity, but also to indicate the relative contributions of each substrate. The latter part will be examined in the section on substrate switch below.

Maximal oxygen consumption rate in the coupled state (CI+ProDH+CII+mtG3PDH-OXPHOS) increased from 19°C to 30-38°C, with slight, non-significant decreases between 34 and 38°C in all species except the heat-tolerant *D. virilis* and *D. mojavensis* (Figs. 2A-C, S2A-C). At 42°C the maximal rates were mostly higher than those measured at 19°C (however only significantly in *D. mojavensis*, Fig. 2C), but lower than at 38°C (though only significantly in *D. virilis*). At the most extreme temperature, 46°C, OCRs in *D. immigrans*, *D. mercatorum* and *D. virilis* were lower compared to the other “high” temperatures (everything above 19°C), but similar to the rates measured at 19°C (Fig. S2A-C), while in *D. subobscura* and *D. melanogaster* the rates at 46°C were significantly lower than at all other temperatures (Fig. 2A,B). The most heat tolerant species, *D. mojavensis*, displayed a different response to the extreme temperature and maintained a high maximal OCR, which was similar to the other “high” temperatures (> 19°C) and significantly higher than the rate measured at 19°C (Fig. 2C). Hence, all species were able to maintain high maximal respiration rates at species-specific temperatures that are incompatible with survival for more than a few minutes.

After the maximal coupled respiration rate had been measured, FCCP was added to measure OCR in the non-coupled state, as this uncoupler allows protons to cross the inner mitochondrial membrane and thus alleviates any limitations to respiration by phosphorylation. The non-coupled respiration rates (FCCP-ETS) are presented in Figs. 2A-C and S2A-C, but another way to examine the potential constraints of the phosphorylating system on the ETS capacity is to calculate the non-coupled ratio (*ETS_max_/OXPHOS_max_*) by dividing the non-coupled rate with the maximal coupled rate (FCCP-ETS/CI+ProDH+CII+mtG3PDH-OXPHOS, Table S2). In all six species, the highest values of *ETS_max_/OXPHOS_max_* were observed at the lower range of temperatures indicating that ETS capacity was not fully coupled to phosphorylation here. *ETS_max_/OXPHOS_max_* decreased with temperature indicating that the full ETS capacity is utilized for phosphorylation at high temperatures (*ETS_max_/OXPHOS_max_* was even frequently below 1 which is the lowest theoretical value of *ETS_max_/OXPHOS_max_* since it indicates that the OCR remained the same following FCCP injection).

Maximal activity of complex IV (CIV), the last respiratory enzyme of the ETS where oxygen is used as the final electron acceptor, was measured after inhibition of complexes I, II and III and was stimulated by the artificial substrate TMPD (along with ascorbate) (Fig. S4). When temperature was increased from 19 to 30°C, all species showed increased CIV activity, which for *D. subobscura*, *D. mercatorum*, *D. virilis* and *D. mojavensis* were significantly higher (*p* ≤ 0.003, one-way ANOVA with Tukey’s *post hoc* adjustment), but did not reach the level of significance in *D. immigrans* and *D. melanogaster* (*p* = 0.121 and 0.696, respectively). At 34°C CIV activity was mostly similar or not significantly increased compared to 30°C (*p* ≥ 0.081), and likewise when CIV activity measured at 38°C was compared to 34°C. However, *D. immigrans* showed a significant increase in CIV activity (*p* = 0.003), and also reached its maximal capacity at this temperature (38°C). *D. mercatorum* and *D. melanogaster* also peaked in measured CIV capacity at 38°C, while *D. subobscura* and *D. virilis* showed a plateau-like CIV capacity from 30 to 38°C, with slightly higher capacities at the lower temperatures. The most heat tolerant species, *D. mojavensis*, peaked at 42°C. This was however not statistically different from the activity observed at 38°C (*p* = 0.993). For the other species, 42°C decreased CIV capacity, although only significantly in *D. immigrans* and *D. mercatorum* (*p* < 0.001 and *p* = 0.007, respectively). At the extreme 46°C, all species showed significantly reduced rates compared to the other “high temperatures”, while the CIV activity was mostly similar to the rate measured at 19°C (only *D. subobscura* showed a significantly lower rate than at 19°C).

### High temperature diminishes mitochondrial coupling at the level of complex I

The OXPHOS coupling efficiency (*j≈p*, unitless) at the level of complex I was calculated as [1-(CI-LEAK/CI-OXPHOS)], a linearized form of the traditional respiratory control ratio (RCR) which describes the flux control of ADP on CI-supported respiration (Table 1).Here *j≈p* approaching 1 indicates a maximally coupled complex I and *j≈p* = 0 indicates a non-ADP controlled complex I respiration. All species showed high values of *j≈p* at 19, 30 and 34°C (range: 0.892 – 0.972, which is above the 0.8 value that is traditionally expected from “healthy”, functional mitochondria (RCR of 5 transformed to *j≈p*) (Gnaiger, 2014)) and within species these values were not significantly different (*p* ≥ 0.949, one-way ANOVA with Tukey’s *post hoc* adjustment). At 38°C *D. subobscura* displayed a reduced *j≈p* (0.384 ± 0.104) compared to the lower temperatures (*p* < 0.003), and similar reductions were found at 42°C for *D. immigrans* (0.195 ± 0.069, *p* < 0.001), *D. mercatorum* (0.381 ± 0.072, *p* < 0.001), *D. melanogaster* (0.252 ± 0.123, *p* < 0.001) and *D. virilis* (0.431 ± 0.136, *p* < 0.006). For *D. mojavensis j≈p* at 42°C was reduced compared to 19 – 34°C (0.377 ± 0.201, *p* <0.009), but the value was not significantly different from 38°C (*p* = 0.058). However, at the most extreme temperature, 46°C, all species displayed highly reduced values of *j≈p* compared to the lower temperatures (Table 1).

### Temperature-dependent shift in mitochondrial substrate oxidation

Using the sequential injection of substrates in the SUIT protocol allowed us to calculate the relative contribution to the OCR for each substrate (the substrate contribution ratio - SCR, Figs. 2D-F, S2D-F). At the lower temperatures (19-34°C), it is clear from the SCRs of all species that addition of the three substrates proline, succinate and glycerol-3-phosphate (G3P) did not stimulate the OCR markedly above that measured with only pyruvate and malate. This is in marked contrast to the situation at higher temperatures where SCRs were elevated for proline and succinate particularly at 42 and 46°C, and SCRs for G3P were very high at 38-46°C. Across species the relative effect of adding proline was smaller than the effect of adding succinate or G3P. Only in *D. subobscura* and *D. mercatorum* was it possible to detect a temperature effect on the stimulation from proline injection and discern between temperatures (with larger responses at higher temperatures) (Figs. 2D, S2E). Succinate gave high values of SCR in *D. immigrans*, *D. subobscura* and intermediate values in *D. melanogaster* at both 42 and 46°C, while large effects of succinate were only found at 46°C in *D. mercatorum*, *D. virilis* and *D. mojavensis*. For G3P, *D. immigrans* (Fig. S2D) and *D. subobscura* (Fig. 2D) showed significant increases in SCR at 38°C, while this effect was only significant at 42°C for the remaining four species (Figs. 2B,C and S2B,C). At 46°C, SCR for G3P decreased in most species except *D. virilis* and *D. mojavensis*, in which the SCR increased (though only significantly in *D. mojavensis* (Fig. 2C)).

The high SCR values indicate that the OCR is markedly stimulated by the injection of these substrates. In other words, we find that when complex I fails (see above), other substrates can take over to support respiration. An additional set of experiments were performed to examine if the increased effect of injection of alternative substrates (proline, succinate and G3P) at high temperatures was attributable to the removal of a complex I “masking effect” through the temperature-induced breakdown of CI-OXPHOS, or rather that the utilisation processes of alternative substrates were temperature-dependent. Specifically, a second SUIT protocol was designed to investigate if proline and G3P could facilitate high OCRs at 34 °C, a temperature where CI-OXPHOS is high (using the standard SUIT protocol, Fig. 2A-C), when their effect on OCR is marginal using the standard protocol at this temperature. The OCRs measured with the two SUIT protocols are not directly comparable, but when we tested *D. subobscura*, *D. melanogaster* and *D. mojavensis*, we found that proline and particularly G3P can sustain high OCRs at 34°C, when complex I is artificially inhibited with rotenone prior to substrate injections (Fig. S5). We also inhibited complex I at 42°C, a temperature where complex I is already markedly depressed when measured using the standard protocol (Fig. 2A-C) and saw a similar large contribution of particularly G3P to oxygen consumption (Fig. S5).

### Breakdown of complex I mediated respiration is not related to a loss of complex I enzyme activity

To examine whether the breakdown of CI-OXPHOS observed at the higher assay temperatures was related to temperature-induced disruption of enzymatic function in the electron transporting enzyme itself, enzymatic catalytic capacities were measured at a range of temperatures (23.5-45°C).

Complex I showed stable increases in enzymatic catalytic capacity with temperature in all species (Figs. 3A, S3A), with no apparent breakdown in contrast to the high-resolution respirometry experiments. Next, we measured pyruvate dehydrogenase (PDH) which oxidizes pyruvate into acetyl-CoA and thus links glycolysis with the tricarboxylic acid cycle while producing NADH that will feed electrons to complex I (Fig. 1). For all species the enzymatic activity of PDH increased with temperature, until conversion rates dropped in species with low to moderate heat tolerance at 42-45°C (*D. immigrans*, *D. subobscura*, *D. melanogaster* and *D. mercatorum*), although not significantly in the latter species (Figs. 3B, S3B). In the heat tolerant species (*D. virilis* and *D. mojavensis*) PDH activity increased or remained the same (as at 42°C) at 45°C. PDH thus showed species-specific responses to increased temperature that may relate to species heat tolerance.

**Fig. 3.**
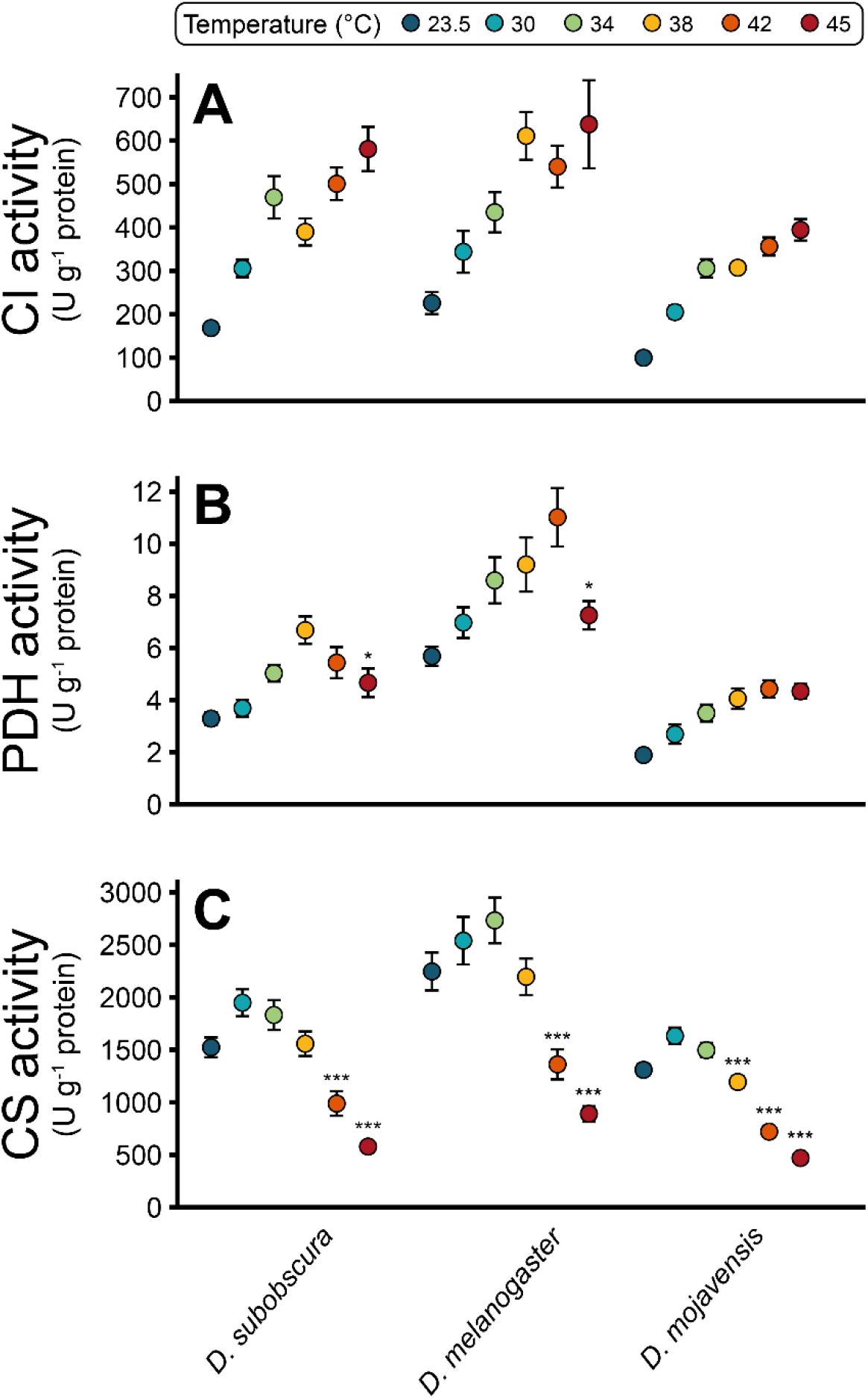
Mitochondrial enzymes display divergent responses to high temperature. Enzymatic activities were measured in homogenized thoraces for A) NADH:ubiquinone oxidoreductase (complex I, CI), B) pyruvate dehydrogenase (PDH) and C) citrate synthase (CS), here shown for three species (see Fig. S3 for the other species). All enzymatic activities are reported as U g^−1^ protein mean ± s.e.m., where U represents 1 μmol substrate transformed to product in 1 minute. Notice that the temperatures used for measurements are not the same as used for high-resolution respirometry (23.5 and 45°C substituted 19 and 46°C, respectively, see Materials and methods). Asterisks denote statistically significant differences between the measurements at higher temperatures than the temperature where the maximal enzymatic catalytic capacity was observed and the maximal rate; *p < 0.05, ** p < 0.01 and *** p < 0.001 in one-way ANOVAs with Tukey post hoc adjustments.

Finally, we measured the activity of citrate synthase (CS) which facilitates the condensation of acetyl-CoA with oxaloacetate to form citrate in the tricarboxylic acid cycle (Fig. 1). CS showed similar reactions to high temperature in all species (Figs. 3C, S3C), with activity increasing from 23.5 to 30°C. Conversion rates levelled at 34°C (with small increases in *D. immigrans* and *D. melanogaster*), then dropped significantly at 38°C (*D. immigrans*, *D. mercatorum* and *D. mojavensis*) and 42°C in *D. subobscura*, *D. melanogaster* and *D. virilis*. Accordingly, citrate synthase displayed heat-induced perturbation of enzymatic catalytic capacity in all six species, however the pattern was not following species heat tolerance to the same degree as observed for PDH activity.

## Discussion

In the present study we measured mitochondrial respiration rates in six *Drosophila* species with differing heat tolerance at temperatures ranging from benign to temperatures around and above species tolerance limits. With this design we examined how temperature affects mitochondrial function, and if loss of mitochondrial function can be related to organismal heat tolerance.

### Mitochondrial function persists at temperatures above species heat tolerance

Several studies on ectotherms suggest that mitochondria are more heat tolerant than the animal as a whole (Chung and Schulte, 2020), and in insects a mitochondrial ‘hyperthermic overdrive’ in which mitochondria perform rapid aerobic metabolism is observed after the loss of higher organismal function (Heinrich et al., 2017; Mölich et al., 2013). In accordance, a recent study on the honey bee *Apis mellifera* found that mitochondrial respiration was intact at 50°C (Syromyatnikov et al., 2019), which is higher or equal to the CT_max_ measured in two subspecies of *A. mellifera* using thermolimit respirometry (Kovac et al., 2014). In the *Drosophila* system we find similar evidence for sustained mitochondrial function at high temperatures since all species were able to maintain high oxygen consumption rates at temperatures above their organismal heat limit (as characterized in Jørgensen et al. (2019)). However, our results show that this mitochondrial heat tolerance is highly dependent on the oxidative substrates used to fuel respiration.

### Hyperthermic breakdown of complex I-supported respiration

Energetic demand increases with temperature in ectotherms, and accordingly the oxygen consumption related to mitochondrial aerobic ATP production must follow. When temperature was increased from the acclimation temperature (19°C) to 30°C, a high yet benign temperature change for all of the tested species, we observed a general increase in maximal oxygen consumption rate under OXPHOS conditions (CI+ProDH+CII+mtG3PDH-OXPHOS), which was primarily driven by increased complex I-supported oxygen consumption (Figs. 2A-C and S2A-C). The OCRs measured in *D. melanogaster* for each step of the SUIT protocol at 19 and 30°C were similar to previous measurements using the same protocol at 24°C in that species (Cormier et al., 2019). At 34°C, however, maximal oxygen consumption stagnated, and in the three least heat tolerant species (*D. immigrans*, *D. subobscura* and *D. mercatorum*), the OCR did not stabilize in some preparations following ADP injection (CI-OXPHOS, Figs. 2, S1,2), which was also observed at 38°C in the same species. Likewise, at 38°C some of the preparations from the heat tolerant species *D. virilis* and *D. mojavensis* displayed this ineptness to maintain a stable CI-OXPHOS, and for these tolerant species the phenomenon persisted at 42°C (Figs. 2, S1,2). It was only in *D. melanogaster*, a moderately heat tolerant species for which this SUIT protocol was optimized, that we did not observe this. Instead this species was characterised by an abrupt decrease in CI-OXPHOS when temperature was increased from 34 to 38°C (Fig. 2B) which was also observed in the stable CI-OXPHOS traces in the other species (at 34-38°C or 38-42°C depending on species). These findings indicate a hyperthermic breakdown of complex I-supported respiration. Indeed, it has been shown that NADH-dependent (i.e. CI-supported) OCRs measured in *in vivo* heat-treated blowflies was reduced by 50 % compared to non-heated controls (El-Wadawi and Bowler, 1996). Complex I has also been suggested to be a primary site of heat failure in liver mitochondria from marine fishes (Chung et al., 2018; Martinez et al., 2016), in marine crustaceans (Iftikar et al., 2010), as well as in maize (Pobezhimova et al., 1996). In the present study the OXPHOS coupling efficiency *j≈P* (the linearized form of the respiratory control ratio, RCR) at the level of complex I, decreased with higher temperature (Table 1), which is an obvious consequence of the hyperthermic decrease in CI-OXPHOS rather than an increase in CI-LEAK, as previously observed (Hilton et al., 2010; Iftikar et al., 2010; Iftikar et al., 2014; Lemieux et al., 2010b). Accordingly, indications of a hyperthermic breakdown of complex I calls for examination of the underlying cause(s) as well as how the mitochondria function with this impairment considering the persistent ability to maintain high maximal OCRs across a wide range of high temperatures.

Complex I, or NADH:ubiquinone oxidoreductase, is the major entry point for electrons into the ETS situated in the inner mitochondrial membrane (Fig. 1) via oxidation of mitochondrial NADH produced by various metabolic pathways such as the TCA cycle, pyruvate oxidation and *β*-oxidation of fatty acids (Hirst, 2010). As the mitochondrial membranes are impermeable to NADH and NAD+, complex I is also an important regulator of the matrix redox pool (NAD+/NADH ratio) which is required for the TCA cycle and various enzyme functions to continue (Sacktor, 1975). Since CI-OXPHOS decreased at high temperatures in all of the species tested, we measured the enzymatic activity of complex I to examine if this decline in activity was due to impaired enzyme function. We found that, at least in homogenised thoraces, complex I activity did not suffer from a hyperthermic breakdown. Instead, enzymatic catalytic capacities increased with temperature, and peaked at 45°C (the highest temperature tested) (Figs. 3A, S3A). Thus, it is likely that the limitation of CI-OXPHOS observed here at high temperature occurs upstream of complex I. A previous study in tobacco hornworm (*Manduca sexta*) reported that the substrate oxidation system governs a significant portion of the temperature effect on maximal mitochondrial respiration (Chamberlin, 2004). The pyruvate dehydrogenase complex (PDH) is a potential candidate explaining the observed decrease in CI-OXPHOS (Blier et al., 2014; Lemieux et al., 2010a) (Fig. 1). PDH activity increased with temperature (as would be expected), but above 38°C, there were species-specific differences in the reaction patterns (Figs. 3B, S3B). In the least heat tolerant species (*D. immigrans* and *D. subobscura*) activity decreased at 42 and 45°C, while in the moderately heat tolerant species (*D. mercatorum* and *D. melanogaster*) it increased up to 42°C and then decreased at 45°C. Finally, in the most heat tolerant species, *D. virilis* and *D. mojavensis*, PDH activity increased continuously or stagnated at 45°C. These interspecific patterns could be related to species heat tolerance, as observed for other enzymes (Dahlhoff and Somero, 1993; Hochachka and Somero, 2002). However, the temperatures at which declines in PDH activity are observed are higher than the temperature interval (34-42°C) where CI-OXPHOS rates were found to decrease, and notably for *D. virilis and D. mojavensis* in which PDH activities do not appear to be compromised at all. Lastly, we also measured the activity of citrate synthase (CS, Fig. 1). Generally CS activities increased from 19 to 30°C but decreased slightly in most species at 34°C before they progressively dropped as temperature was raised to 45°C (Figs. 3C, S3C). Unlike for PDH activity, there was no clear pattern for the decrease in enzymatic activity between species that could potentially be related to their heat tolerance. Instead, it seems that CS is generally challenged at high temperatures in *Drosophila*. Heart CS (and PDH) activity was found to decrease at temperatures around and exceeding CT_max_ in European perch (*Perca fluviatilis*), which was interpreted as an impaired capacity to oxidize pyruvate which could ultimately limit the entry of electrons into the ETS (Ekström et al., 2017). However, the literature is ambiguous on the potential limitation of CS (and PDH) on ectotherm metabolism at high temperatures and it has been disputed in mussels (Hraoui et al., 2020). Nevertheless, it must be noted that the enzymatic activities measured *in vitro* here represent the maximal catalytic capacity and that they may not directly reflect metabolic flux *in vivo*. Thus, the temperature mismatch between enzymatic activities and CI-OXPHOS breakdown could suggest involvement of other components of the substrate oxidation system, e.g. the mitochondrial pyruvate carrier. Although the decreased catalytic capacities of CS and PDH activities may create a bottleneck for NADH production by the TCA cycle, this cannot fully explain the drastic drop in CI-OXPHOS observed at high temperatures. A possible explanation would be that allosteric regulation and/or covalent modifications occur at the level of complex I at high temperature, reducing its ability to oxidize NADH. The present study shows that complex I-supported oxygen consumption is challenged at high temperatures across the tested *Drosophila* system, with abrupt declines in CI-OXPHOS at temperatures that tend to relate to the species heat tolerance, but also that this decline is not likely to be an effect of heat perturbations on the complex I enzyme itself. Instead the enzymes PDH and CS which are working downstream of the ETS showed decreased activities at high temperatures and may thus point to a cause of the observed decrease in CI-OXPHOS; that the substrate oxidation system fails to provide the appropriate amount of NADH to complex I.

### Mitochondrial flexibility: oxidation of alternative substrates sustains high maximal oxygen consumption rates

Insect species have different preferred substrate for flight metabolism; short-term fliers like flies and bees (*Diptera* and *Hymenoptera*) mainly use carbohydrates while locusts and butterflies (*Orthoptera* and *Lepidoptera*) use fats as fuel for sustained flight (Chadwick and Gilmour, 1940; Krogh and Weis-Fogh, 1951; Sacktor, 1955). A survey of the literature on insect mitochondrial respiration presented by Soares et al (2015) indicated that *Drosophila* are particularly relying on NADH-dependent (i.e. electron donors for complex I) respiration, which includes pyruvate, malate and proline. Notice that proline is included as an electron donor for complex I as it can be transformed into α-ketoglutarate and thus increase TCA cycle intermediates to oxidize pyruvate (anaplerotic role, Fig. 1) (Sacktor and Childress, 1967). However, proline is also recognized as a direct electron donor to the ubiquinone pool through the flavoenzyme proline dehydrogenase (Bursell, 1981; McDonald et al., 2018; Olembo and Pearson, 1982; Soares et al., 2015; Teulier et al., 2016), and some insects are even relying on proline as their main fuel (Bursell, 1963; Teulier et al., 2016). G3P is also an important oxidative substrate for insects allowing the entry of electrons into the ETS via the mtG3PDH, as it is at the intersection of glycolysis, fatty acid degradation and oxidative phosphorylation (McDonald et al., 2018). In insect flight muscle, this reaction is the most important redox cycler (glycerol phosphate shuttle) for maintaining redox balance (NAD+/NADH) in the cytosol (Sacktor, 1975), attested by *Drosophila* mutants with reduced or absent mtG3PDH displaying debilitated flight ability (Carmon and MacIntyre, 2010; Carmon et al., 2010). However, almost nothing is known about the regulatory mechanisms at the three mitochondrial loci of metabolic control in insects during temperature changes; namely, the oxidation of pyruvate, proline and G3P.

The oxygen consumption rates measured in the present study on *Drosophila* at benign temperatures also indicated that pyruvate (as NADH-dependent or CI-OXHOS) was the most efficient fuel for the ETS, congruent with the data compiled by Soares et al (2015). Once temperature increased and the breakdown in CI-OXPHOS was observed, the substrate contribution ratio (SCR) of the other provided substrates (proline, succinate and glycerol-3-phosphate (G3P)) increased (Figs. 2, S2), but this is not surprising as the calculation of SCR is based on the OCR achieved by the previous substrate. Hence, we investigated if the “newfound” use of alternative substrates with increasing temperature was solely due to the loss of CI-OXPHOS (here labelled the *masking effect*), or if the pathways for the alternative substrates were increasingly active due to the higher temperatures. To examine this, we conducted a second set of measurements on permeabilized thoraces with a new SUIT protocol, inhibiting complex I with rotenone prior to injection of alternative substrates (proline and G3P) at 34°C (when CI-OXPHOS is not usually compromised) and 42°C (when CI-OXPHOS is “naturally” reduced). We found that proline and particularly G3P efficiently maintained high rates of oxygen consumption (Fig. S5). Together with the increased values of SCR for proline and G3P at high temperatures, this indicates that a *masking effect* from complex I dominates the apparent contribution of alternative substrates to the OCR, rather than a clear temperature effect on alternative pathways.

To summarise, when CI-OXPHOS is challenged by high temperatures, other substrates can be oxidized instead to deliver electrons to the ETS. This switch in substrate maintains a high maximal oxygen consumption rate even at temperatures above those normally considered lethal. In model simulations of mitochondrial flexibility in human cardiomyocyte mitochondria with complex I deficiency, Zieliński et al. (2016) identified two dominant mechanisms to maintain redox balance. These reactions were the glycerol phosphate shuttle (cytosolic) and the cycle between proline dehydrogenase and pyrroline-5-carboxylate reductase (mitochondrial). In accordance, it has been shown that reduced pyruvate-supported oxygen consumption in mutant *Drosophila* is associated with a compensating increase by proline dehydrogenase to mitochondrial respiration (Simard et al., 2020a; Simard et al., 2020b). Similarly, decreased mitochondrial respiration at the level of complex I in mutant *Drosophila* was associated with an increase in the mitochondrial G3P oxidation, compensating for complex I deficiency (Pichaud et al., 2019). Our data suggest that the same reactions are important when CI-OXPHOS is reduced by heat stress in *Drosophila*.

In the model simulations from Zieliński et al. (2016) the ATP production was somewhat reduced when G3P and proline were used as alternative substrates since these reactions do not contribute directly to the proton gradient (only indirectly through complex III and IV through the downstream ubiquinone pool). Considering the failure of complex I to support the proton gradient at high temperature, it is possible that the sustained oxygen consumption rates at high temperatures are somewhat decoupled to sustained ATP production rates in *Drosophila*. As an example it was found in the wrasse *Notolabrus celidotus* that respiration through complex I and II in permeabilized cardiac fibres increased at temperatures above the upper tolerance limit but that the concurrent ATP production rate plummeted resulting in a lower ATP/O ratio (Iftikar and Hickey, 2013; see also Chung and Schulte, 2020). The decreased ATP/O coincides with the temperature of acute heart failure, emphasizing that sustained high mitochondrial oxygen consumption rates may not necessarily result in efficient energy production. Interestingly, mutants of *D. subobscura* with reduced activity of complex I (−50%) and complex III (−30%) compared to the wildtype showed unaltered ATP production, and the authors further found that complex I activity could be inhibited up to 70% (in wildtype) before any change was observed in OCR and ATP synthesis (Farge et al., 2002). Further studies should therefore investigate if the characteristic hyperthermic failure of CI-OXPHOS result in insufficient ATP production or if energy production is well defended at critically high temperatures.

Heat stress also increases the production of reactive oxygen species (ROS), which are normal by-products of cellular respiration and represent both an important signaling molecule and a potential source of cellular injury (Abele et al., 2002; Blier et al., 2014; Scialò et al., 2020). For example, a recent paper showed that limiting ROS produced in brain mitochondria by reverse electron transport through complex I (ROS-RET) during heat stress in *D. melanogaster* reduced survival, as ROS-RET activates survival responses (Scialò et al., 2020). On the other hand heat stress has been found to increase ROS production (Abele et al., 2002; Lopez-Martinez et al., 2008; Wang et al., 2014; Yang et al., 2010; Zhang et al., 2015). Complex I is normally considered the main source of ROS (Murphy, 2009) but in *Drosophila*, mtG3PDH is one of the most significant producers of ROS (Miwa and Brand, 2005; Miwa et al., 2003). Investigating the ROS production at high temperatures is thus an interesting direction for future studies given the breakdown in CI-OXPHOS, the resulting increase in relative contribution of mtG3PDH and the sustained high mitochondrial oxygen consumption rates observed here.

In summary, our study shows that mitochondrial oxygen consumption is sustained at temperatures around and above the species heat tolerance limits, indicating that mitochondria may be more tolerant than the animal itself, in accordance with previous studies. In all of the tested species we observed abrupt declines in the OCR supported by complex I substrates, in a pattern that tends to correlate with species heat tolerance. However, the resolution in assay temperatures is not sufficient to fully discern whether organismal heat tolerance dictates the temperature of CI-OXPHOS collapse in *Drosophila*. Measurements of enzymatic activity revealed that the complex I enzyme alone is unlikely to be responsible for the observed decline in CI-OXPHOS. Instead, upstream enzymes of the substrate oxidation system like the pyruvate dehydrogenase complex and citrate synthase may potentially limit CI-OXPHOS at high temperatures due to their decline in activity. Using an alternative SUIT protocol we also observed that the increased relative reliance on G3P and proline are largely attributable to a *masking effect* of complex I-supported respiration.

Hence, mitochondrial oxygen consumption persists even at critically high temperatures through the use of alternative substrates. It is, however, unclear whether this oxygen use is coupled to sufficient energy production. Future studies should include examination of the ATP production at high temperature to investigate whether the high mitochondrial oxygen consumption is coupled to energy production, and further if the oxygen consumption in combination with alternative substrate pathways and high temperature can result in increased ROS production, ultimately posing an oxidative stress challenge in *Drosophila* near their upper thermal limit.

## Abbreviations

acetyl CoA: acetyl Coenzyme A
ADP: adenosine diphosphate
ATP: adenosine triphosphate
asc: ascorbate
cG3PDH: cytoplasmic glycerol-3-phosphate dehydrogenase
CI: complex I (NADH:ubiquinone oxidoreductase)
CII: complex II (succinate dehydrogenase)
CIII: complex III (coenzyme Q:cytochrome *c* oxidoreductase)
CIV: complex IV (cytochrome *c* oxidase)
CS: citrate synthase
CT_max_: Critical thermal maximum
CV: complex V (ATP synthase)
Cyt *c*: cytochrome *c*
DHAP: dihydroxyacetone phosphate
ETS: electron transport system
ETS_max_/OXPHOS_max_: non-coupled ratio
FADH_2_: flavin adenine dinucleotide
FCCP: carbonyl cyanide-4-(trifluoromethoxy)phenylhydrazone
G3P: glycerol-3-phosphate
GAP: glyceraldehyde-3-phosphate
*j≈P*: OXPHOS coupling efficiency
LEAK: non-coupled (to phosphorylation) respiration
mtG3PDH: mitochondrial glycerol-3-phosphate dehydrogenase
MPC: mitochondrial pyruvate carrier
NADH: nicotinamide adenine dinucleotide
OCLTT: oxygen- and capacity-limited thermal tolerance
OXPHOS: oxidative phosphorylation
PDH: pyruvate dehydrogenase
ProDH: proline dehydrogenase
Q: ubiquinone pool
ROX: residual oxygen consumption
saz: sodium azide
SCR: substrate contribution ratio
TMPD: (N,N,N’,N’-tetramethyl-p-phenylenediamine)

## Acknowledgements

The authors would like to thank Rebekah Strang and Kirsten Kromand for animal care in Canada and Denmark, respectively, Professors Angela Fago and Tobias Wang for allowing us to use their Oroboros systems at Aarhus University, and Dr. Amanda Bundgaard for valuable discussions.

